# Insights into impact of DNA copy number alteration and methylation on the proteogenomic landscape of human ovarian cancer via a multi-omics integrative analysis

**DOI:** 10.1101/488833

**Authors:** Xiaoyu Song, Jiayi Ji, Kevin J. Gleason, John A. Martignetti, Lin S. Chen, Pei Wang

## Abstract

In this work, we propose iProFun, an ***i***ntegrative analysis tool to screen for ***Pro***teogenomic ***Fun***ctional traits perturbed by DNA copy number alterations (CNA) and DNA methylations. The goal is to characterize functional consequences of DNA copy number and methylation alterations in tumors and to facilitate screening for cancer drivers contributing to tumor initiation and progression. Specifically, we consider three functional molecular quantitative traits: mRNA expression levels, global protein abundances, and phosphoprotein abundances. We aim to identify those genes whose CNAs and/or DNA methylations have cis-associations with either some or all three types of molecular traits. In comparison with analyzing each molecular trait separately, the joint modeling of multi-omics data enjoys several benefits: iProFun experienced enhanced power for detecting significant cis-associations shared across different omics data types; and it also achieved better accuracy in inferring cis-associations unique to certain type(s) of molecular trait(s). For example, unique associations of CNA/methylations to global/phospho protein abundances may imply post-translational regulations.

We applied iProFun to ovarian high-grade serous carcinoma tumor data from The Cancer Genome Atlas and Clinical Proteomic Tumor Analysis Consortium, and identified CNAs and methylations of 500 and 122 genes, respectively, affecting the cis-functional molecular quantitative traits of the corresponding genes. We observed substantial power gain via the joint analysis of iProFun. For example, iProFun identified 130 genes whose CNAs were associated with phosphoprotein abundances by leveraging mRNA expression levels and global protein abundances. By comparison, analyses based on phosphoprotein data alone identified none. A group of these 130 genes clustered in a small region on Chromosome 14q, harboring the known oncogene, *AKT1*. In addition, iProFun identified one gene, *CANX*, whose DNA methylation has a cis-association with its global protein abundances but not its mRNA expression levels These and other genes identified by iProFun could serve as potential drug targets for ovarian cancer.

## INTRODUCTION

The initiation, progression and metastasis of cancer often results from accumulation of DNA-level variations, such as DNA copy number alterations (CNAs) and epigenetic modifications. CNAs involve gains or losses of a large region of tumor DNA that could result in activation of oncogenes or inactivation of tumor suppressors, respectively [1, 2, 3]. Hypermethylation of CpG islands often results in silencing the expression of DNA repair genes when occurring in the promoter region and activating oncogenes if in the coding regions [4, 5, 6, 7, 8]. Most major cancer types have been systematically profiled for copy number and CpG methylation, and as a result many CNAs and DNA methylations have been identified and been associated to carcinogenesis and cancer progression [9, 10, 11, 12, 13, 14]. However, it remains challenging to pinpoint diagnostic, prognostic and therapeutic targets from this long list of cancer-associated genes. In particular, it is important to distinguish the driver genes that contribute to oncogenesis and cancer progression from the passengers acquired by random alterations during cancer evolution [2] and changes in gene activities that are the consequences, not causes, of cancer. To address this challenge, previous studies have primarily focused on associating CNAs and DNA methylations to their cis (i.e., local) gene expression levels [15], a form of molecular quantitative trait (QT) that is relatively easier to measure than protein abundances. Significant associations of CNAs and methylations with cis-mRNA expression levels partially reveal the molecular mechanisms of cancer-associated genes.

In addition to mRNA expression levels, it is of great interest to characterize the functional consequences of CNAs and DNA methylations on protein abundances. Proteins and phosphoproteins are key molecules that carry out most cellular functions, and are essential to cancer initiation, tumor progression and response to therapy. The observed median correlations between mRNA gene expressions and global proteomic abundances, when measured and quantified in tumor tissues, are 0.45, 0.39 and 0.47 in ovarian [16], breast [17] and colorectal tumors [18], respectively. In addition to global protein abundances, phosphorylation often occurs on multiple distinct sites of a given protein, facilitating complex, multi-level regulation that is not reflected at the mRNA expression levels. By investigating the effects of CNAs [19] and separately the effects of DNA methylations [20] on global and phospho proteomic changes, novel insights have been obtained. However, due to a lack of proper analytic tools, few studies have employed an integrative approach to evaluate the impact of CNAs and DNA methylation on multi-level molecular QTs in tumor genomes from a systematic perspective.

Motivated by these challenges and needs, in this work we propose to conduct integrative analysis of multiple types of omics data in order to achieve a systematic and comprehensive understanding of the functional mechanisms of DNA-level alterations in tumors. We propose a novel ***i***ntegrative analysis tool to screen for ***Pro***teogenomic ***Fun***ctional traits altered by CNA and DNA methylation (iProFun). Specifically, we are interested in (1) detecting genes with “cascading effects” on downstream molecular traits, i.e., a gene’s DNA alterations have associations with its cis mRNA expression levels, and global and phospho protein abundances; and (2) identifying associations unique to certain type(s) of molecular trait(s), in particular unique to global/phospho protein levels. The associations of DNA alterations unique to the protein levels but not reflected on mRNA levels suggested post-translational regulation. Despite the urgent needs for integrative analysis methods and tools in biomedical research, the integration of data from multiple data types also imposes tremendous statistical challenges, such as high dimensionality, complex gene-gene correlations, different scales and distributions among different types of omics data, and complete or partial overlapping of samples across platforms [21, 22]. To address these challenges, the iProFun method takes as input genome-wide summary statistics in assessing associations of CNVs and DNA methylations on each type of molecular traits, and allows genes and molecular QTs to be arbitrarily correlated in the joint analysis. The iProFun method estimates the conditional density of each type of trait separately, allowing different scales and distributions among different data types. And, it also allows for sample correlations due to complete or partial overlapping of samples. In comparison with the separate analyses of CNA and then methylation on each molecular trait, iProFun leverages data from multiple sources and follows information across data types. By imposing rigorous assessment of false discovery rates (FDR), we have demonstrated that iProFun is able to largely boost power and maintain low FDR in identifying various types of cis-associations, in particular in the data types with relatively low sample sizes.

We applied iProFun to the ovarian high-grade serous carcinoma (HGSC) data from TCGA and the genome-wide proteomic data measured in CPTAC. HGSC is the leading cause of gynecologic cancer death in the US [23] that most women will present with advanced-stage disease and ultimately die of their disease within five years [24]. Thus, new treatments and an improved understanding of the biologic basis of this cancer are desperately required. Using iProFun we identified a collection of genes whose molecular functional traits at transcriptomic, proteomic and/or phosphoproteomic levels were altered by somatic CNAs and DNA methylations. Some candidates in this list could serve as potential drug targets.

## METHODS

### TCGA-CPTAC ovarian cancer data

The tumor sample data we analyzed were from 570 adults with ovarian HGSC collected by TCGA. The somatic CNA, mRNA and DNA methylation data were downloaded from the TCGA Firehose pipeline processed in July 2016 at the Broad Institute (http://gdac.broadinstitute.org/). The germline genotyping data were obtained from NCI’s Genomic Data Commons [25], and the proteomic and phosphoproteomic data were obtained from the CPTAC Data Portal [16]. CNAs of 11,859 genes were measured in 559 samples. DNA methylations for a total of 25,762 methylation sites from 14,269 genes were measured using the Illumina 27K platform on 550 samples. The mRNA expression levels of 15,121 genes were measured on 569 samples with the microarray platforms. The extensive MS-based proteomic and phosphoproteomic abundances after preprocessing procedures were provided by CPTAC for a combined total of 7,061 proteins from 174 tumor samples and a total of 10,057 phosphosites from 2,865 phosphoproteins in a subset of 69 tumor samples. The same data were used in The NCI-CPTAC DREAM Proteogenomics Challenge. More information about the samples and the associated metadata is available online [16, 26].

### Integrative analysis pipeline

Figure 1 illustrates the integrative analysis pipeline - iProFun - for revealing dynamic *cis* regulatory patterns in tumors. Briefly, iProFun takes as input the association summary statistics from associating CNAs and methylations of genes to each type of cis-molecular trait, aiming to detect the joint associations of DNA variations and molecular traits in various association patterns. Of particular interest are the genes with “cascading effects” on all cis molecular traits of interest and the genes whose functional regulations are unique at protein/phosphoprotein levels. iProFun can incorporate prior biological knowledge through a filtering procedure, and can identify significant genes with calculated posterior probabilities exceeding a threshold while assessing the empirical FDR (eFDR) through permutation. Downstream enrichment analyses were also embedded into our pipeline to allow for more direct interpretations of different association patterns.

**Figure 1:**
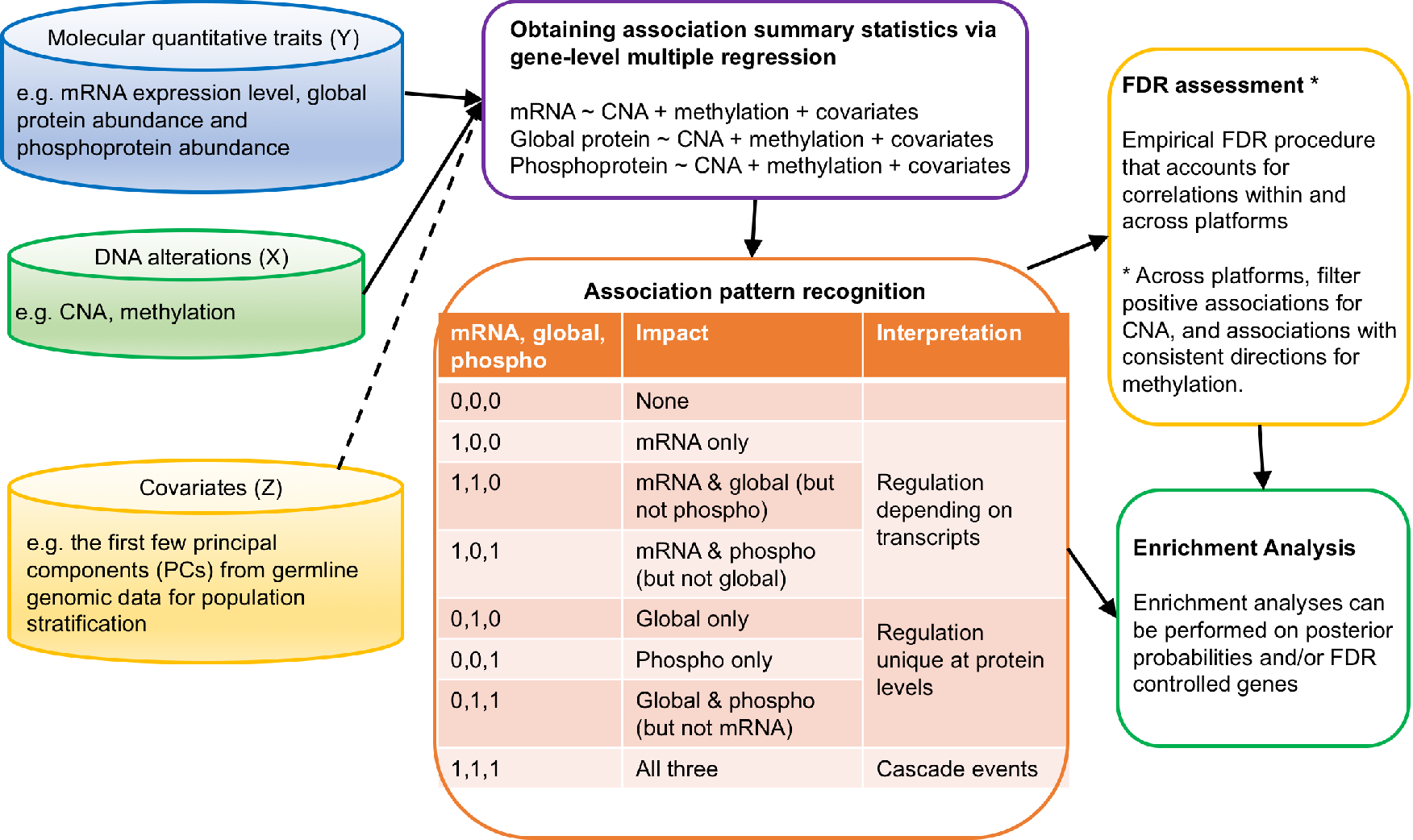
Schematic of iProFun tool for integrative functional analysis

### Preprocessing

All of the five data types were preprocessed to eliminate potential issues for integrative analysis, such as batch effects, missingness and major unmeasured confounding effects. The ovarian HGSC proteomic data were generated at two independent CPTAC centers at Johns Hopkins University and Pacific Northwest National Laboratory. By analyzing the overlapping samples processed in both centers, center effects were corrected using linear normalization to match measured protein abundances. For each of the molecular traits, we filtered genes with missing rate ≥ 50% and then transformed the expression or abundance distributions to standard Gaussian distributions using quantile-quantile normalization. The gene-level CNAs were summarized from all probes nested in the gene by taking their median values. The methylation measures were calculated by beta values ∈ [0,1] with 0 being unmethylated and 1 being fully methylated. To account for potential population structures and other major unmeasured confounding factors, when obtaining the summary statistics for associations, we adjusted for the top Principal Components (PCs) calculated based on germline genotype data, using principal component analysis (PCA) provided in PLINK 1.9. Blood-derived DNA samples were used as the primary source of germline genotype data, with solid normal tissue samples used as a surrogate for subjects who were missing blood-derived DNA samples. We restricted the PCA to bi-allelic variants on autosomes that met the following criteria: minor allele frequency ≥ 0.05; Hardy-Weinberg Equilibrium p-value ≥ 0.0001; pairwise linkage disequilibrium *r*^2^ ≤ 0.2. Finally, we restricted our analysis to genes with quantitative measurements across all five data types (CNA, methylation, mRNA expression, global protein and phosphoprotein) to understand their joint association patterns. A total of 676 genes were analyzed in the following steps.

### Gene-level multiple linear regression to obtain summary statistics

Consider a total of *G* genes that passed the prepossessing procedures in all of the five data types; *n*_1_ samples have measurements of mRNA, CNA and methylation; and *n*_2_ samples have measurements of global protein, CNA and methylation; and *n*_3_ samples have measurements of phosphoprotein, CNA and methylation. We use the following regression models for each type of molecular trait of interest:

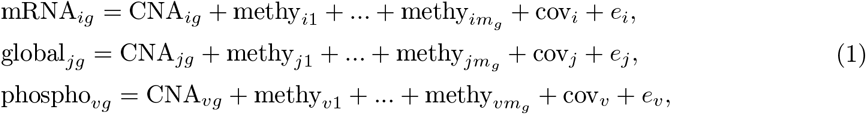

where *i* ∈ (1, …, *n*_1_), *j* ∈ (1, …, *n*_2_) and *v* ∈ (1, …, *n*_3_) are sample indices, *g* ∈ (1, …, *G*) is the index for genes, and cov’s are the sets of covariates adjusted in the regression analyses. The mRNA, global protein and CNA have been summarized at the gene level prior to the analysis, and as such only one variable for each data type is included. Multiple methylation sites may exist for a gene. We use *m_g_* to denote the number of methylation sites for gene *g*, and jointly consider all *m_g_* methylation sites in the regression. Also, multiple phosphosites might exist for a gene. We conducted a pre-screening analysis to select the representing phosphosite from all measured sites of a gene by ANOVA procedure. More specifically, we selected the phosphosite that minimizes the p-value from ANOVA F-test by testing the null model (phospho_*vg*_ = cov_*v*_ + *e_v_*) versus the alternative model (phospho_*vg*_ = CNA_*vg*_ + methy_*v*1_ + … + methy_*vm_g_*_ + cov_*v*_ + *e_v_*). In the above sets of regressions, we adjust for the top three genotype-generated PCs for all three regression models. We adjust for only the top 3 PCs based on a separate investigation in which we compare the regression results adjusting for up to 15 principal components (see Supplementary Figure S1).

We use sets of separate regressions in (1) in the integrative analysis pipeline to allow for different samples being measured on different sets of molecular features. In comparison, a joint analysis of data from all five types may have only a limited number of samples with complete measurements on all of the five data types, while separate analyses of data from subsets of platforms without integration would ignore the potential correlations and connections among different types of genomic features. From this perspective, iProFun is especially appealing in boosting study power for associations with proteomic features, because protein and phosphoprotein data are often measured on much smaller sets of samples than that of mRNA data, and as such power to detect associations with proteins is also much lower. More generally, the integration of summary statistics in iProFun also allows one to take advantage of existing summary statistics when raw data are not available.

### Primo – An integrative analysis method for detecting joint associations of DNA alterations with multi-omics traits

With the summary association statistics obtained from equations (1), we apply an integrative analysis method - Primo - to detect joint associations of DNA variation with multi-omics traits [27, 28].

Consider a total of *G* genes, and *J* sets of summary statistics in assessing the associations of CNAs on *J*-types of cis-molecular traits of interest (here *J* = 3) in (1). Let **T** denote a *G* × *J* matrix of *t*-statistics. When jointly analyzing the association statistics for each gene, there are *K* = 2^*J*^ = 8 possible association patterns. Figure 1 shows the list of all association patterns, as well as the interpretation for each association pattern. Let *q_kj_* = 1 denote a true association between CNA and the *j*th-type of trait under the *k*-th pattern, and *q*_*kj*_ = 0 denote no association. Then (0,0,0) denotes that the CNA has no association with any of the three types of traits, while (0,0,1) and (1,1,1) denote that the CNAs that have effects on only phosphoproteins, and have cascade effects on all of the three traits, respectively.

Consider a total of *G* genes, and *J* sets of summary statistics in assessing the associations of CNAs on *J*-types of cis-molecular traits of interest (here *J* = 3) in (1). Let **T** denote a *G* × *J* matrix of *t*-statistics. When jointly analyzing the association statistics for each gene, there are *K* = 2^*J*^ = 8 possible association patterns. Figure 1 shows the list of all association patterns, as well as the interpretation for each association pattern. Let *q*_*kj*_ = 1 denote a true association between CNA and the *j*th-type of trait under the *k*-th pattern, and *q*_*kj*_ = 0 denote no association. Then (0,0,0) denotes that the CNA has no association with any of the three types of traits, while (0,0,1) and (1,1,1) denote that the CNAs that have effects on only phosphoproteins, and have cascade effects on all of the three traits, respectively.

For each CNA, there must be one and only one true association pattern. Let π_*k*_ denote the “frequency” of CNAs in the *k*-th pattern in the current data. Then the probability of CNA *i* following pattern *k* is given by:

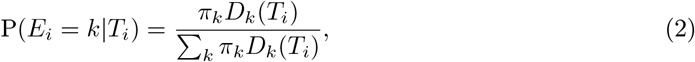

where *D_k_*(·) is the conditional joint density function of the set of *t*-statistics of CNA *i* on *J* traits, conditioning on the *k*-th association pattern. In estimating the conditional joint density, we assume that the *t*-statistics across data types are independent conditional on the current association pattern. That is

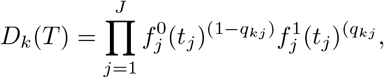

where 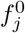
(·) and 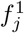(·) represent the marginal null and alternative density functions for the association statistics to the *j*-th trait, respectively. For example, the second pattern (CNA impacts mRNA only) in Figure 1 is (1,0,0), so 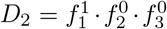 Note that we do not assume statistics from different traits to be marginally independent; instead, we use π_*k*_ to capture both biological correlations among molecular traits and sample correlations among different sets of samples. We estimate the π_*k*_’s for all association patterns in the current data using the Expectation-Maximization (EM) algorithm [29]. Conditional on the data-dependent estimates of π_*k*_’s, the density functions (i.e. distributions) of association statistics on different sets of traits are assumed to be independent. When considering the association patterns as mutually exclusive “clusters”, the conditional joint density can be viewed as the “class centroid.” The estimation of the posterior probability of a CNA following a particular association pattern calculates the relative strength of that pattern in comparison to other patterns.

The Primo method uses the R limma approach [30, 31] to estimate the null and alternative density functions *f*’s for the *J* sets of association statistics. Limma pools information across all genes within each set of statistics, and estimates the empirical nulls as *t*-distributions and the empirical alternatives as scaled *t*-distributions with an estimated scaling parameter. In applying limma and Primo, one needs to specify *a priori* the estimated proportion of non-null statistics (i.e., statistics with non-zero associations) in each set of summary statistics. Those non-null proportion estimates can be obtained from the literature or estimated based on the data. Primo is insensitive to under-specification of the parameters within a reasonable range. In this analysis, we specify that 5% (which is an under-estimation) of association statistics are from the alternative for the effects of CNA on each of the three traits.

Separately, we applied the Primo method to analyze the effects of methylation sites on three types of traits, with similar parameter specifications, and obtained the results for the effects of methylations on cis-molecular traits.

## False discovery rate assessment

In addition to calculating posterior probabilities for each association pattern for each gene, we proposed to also calculate the empirical FDR based on permutations. The empirical FDR can serve as an additional significance measure in accounting for the data-dependent correlation structures and sample overlapping among different data types.

To calculate the empirical FDR, we first calculated the posterior probability of a predictor being associated with an outcome, by summing over all patterns that are consistent with the association of interest. For example, the posterior probability of a CNA being associated with mRNA expression levels was obtained by summing up the posterior probabilities in the following four association patterns - “CNA affecting mRNA only”, “CNA affecting both mRNA & global protein”, “CNA affecting both mRNA & phophoprotein” and “CNA affecting all three traits”, all of which were consistent with CNA being associated with mRNA expression. Then, we permuted our sample 100 times to re-calculate the summary statistics and use them for calculating empirical false discovery rates. More specifically, for each molecular trait, we randomly permute the sample label of the trait while keeping the labels of the other two traits. Then, we re-ran gene-level multiple regressions and Primo analysis and calculated the posterior probability of a predictor being associated with a trait based on the null data sets. Then, for a pre-specified posterior probability cut-off value *α*, a gene was considered positive if its posterior probability is > *α*. We calculated empirical FDR as

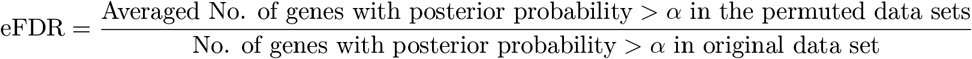

for a pre-specified *α* value. We consider a grid of *α* values, 75% < *α* < 100%. We consider a minimum of 75% posterior probability and an averaged empirical FDR < 10% as the joint significance criteria in declaring significant associations.

Additionally, we added the following filtering procedure to incorporate prior biological knowledge. Literature has suggested that CNA amplifications have often been associated with increased molecular quantities and CNA deletions have been linked with decreased molecular quantities [26, 17, 18]. Meanwhile, variations in methylation might have associations in both directions. For example, hypermethylation in promoter regions may silence DNA repair genes (negative association), but hypermethylation in coding regions may activate oncogenes (positive association). Based on the criteria, only genes with association directions matching biological knowledge (all positive for CNA and same direction for methylation across all traits) were retained and further assessed by empirical FDR.

## RESULTS

### Landscape of CNA and DNA methylation association patterns on functional molecular quantitative traits

To characterize the impact of CNAs and DNA methylations on mRNA, protein and phosphoprotein abundances in ovarian tumors, we applied iProFun on TCGA and CPTAC data as described in Methods. The posterior probabilities of 8 cis-regulation association patterns (Figure 1) of each gene-level CNA and CpG site methylation were calculated (Supplementary Tables S1 and S2). The average posterior probabilities of each cis-regulation association pattern across all CNAs and methylation sites are shown in Figure 2(a). On average, there is a 60% chance for the CNAs to be from the “all three” cis-regulation association pattern, i.e. CNAs are associated with all three cis-QTs: mRNA expression level, global protein and phosphoprotein abundances. The next most common association pattern for CNAs is “mRNA & global” (28%), with “mRNA only” the third most common category (10%). Only 2% of CNAs are estimated to be from the “none” association pattern, i.e. no association between CNAs and any of the three cis-QTs. On the other hand, for 1,103 DNA methylation sites of the 676 genes, on average there is only an 8% chance for the “all three” association pattern, while 52% of the methylation sites have effects on “none” of the traits. Overall, there is a 31% chance for methylation sites to have a cis-regulation with transcripts (18% “mRNA only”, 11% “mRNA & global”, and 2% “mRNA & phospho”), and 8% for unique cis-association with protein levels (0% “global only”, 6% “phoshpo only” and 2% “global & phospho”). It is clear that the effects of CNAs on cis-molecular QTs are much stronger than those of DNA methylations. Enrichment analyses based on posterior probabilities identified that chromosome arms 3p, 8q and 10q are enriched with cascading genes with CNA associations to “all three” cis-molecular traits (Supplementary Figure S2).

**Figure 2:**
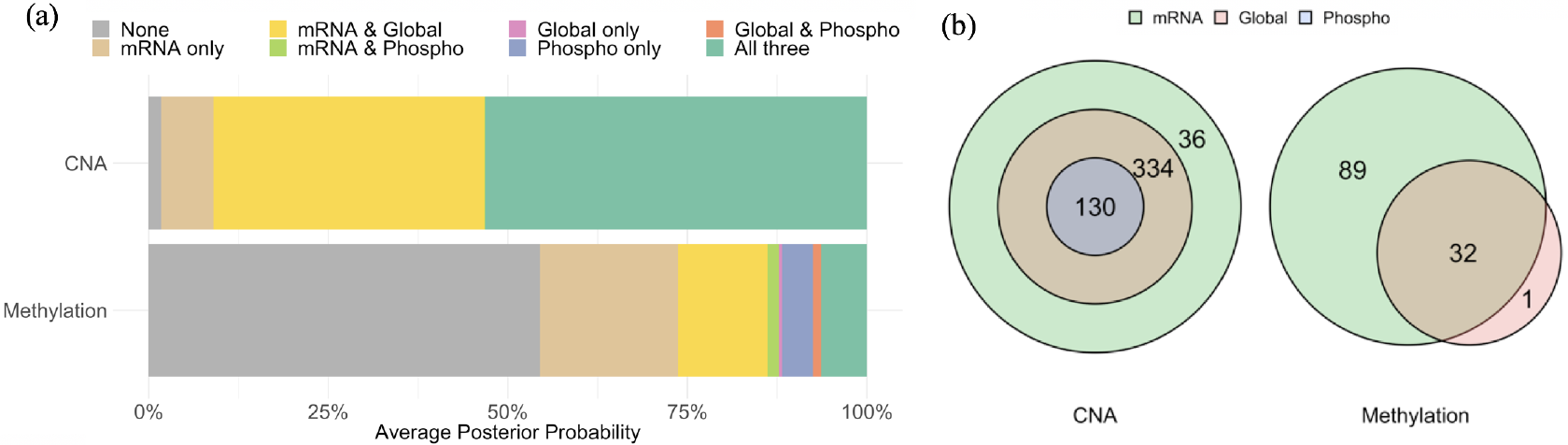
(a) The averaged posterior probabilities of association patterns of CNA and DNA methylation on molecular quantitative traits. (b) The Venn diagram of the identified CNAs and methylations with significant effects on molecular quantitative traits at FDR <10% and posterior probability >75%.

After further deriving the empirical FDRs for all cis-associations based on permutation tests (see Methods), we selected significant association pairs by requiring posterior probabilities >75% and empirical FDRs < 10%. Figure 2(b) displays the Venn diagrams of the numbers of genes whose CNAs and/or DNA methylations were significantly associated with their cis-mRNA levels, global or phosphoprotein abundances. A full list of CNAs and DNA methylation sites with significant cis-associations are provided (Supplementary Tables S3 and S4).

Out of the 676 gene-level CNAs in our analysis, 36 CNAs were associated with mRNA levels only; 334 CNAs were associated with mRNA levels and global protein abundances, but not phosphoprotein abundances; while 130 were identified as *cascade CNAs*, i.e. the CNA of one gene demonstrates significant association with all of the three traits (mRNA levels, protein and phosphoprotein abundances). Network analysis of these 130 genes with cascade CNA cis-associations using the STRING V10 database [32] (Supplementary Figure S5) revealed *AKT1* as a key hub node interacting with many other cascade CNA genes. The gene *AKT1*, located on 14q32.33, is an important effector of the PI3K/RAS pathway. Detailed information of all five omics AKT1 data across the 69 samples are presented in Figure 3. The dosage effects of *AKT1* copy number gains on its gene expression levels, protein and phosphoprotein abundances is clearly demonstrated.

Out of 1103 methylation sites of 676 genes, none revealed significant association with corresponding phosphoprotein abundances. There were 32 methylation sites from 32 genes associated with both their mRNA expression levels and global protein abundances. Interestingly, four out of these 32 genes — *DNMT3A, RHPN2, MAP2*, and *CDH6* — also had cascade CNA cis-associations. The detailed information of all omics data of these four genes are illustrated in Figure 3. On the other hand, the methylation site (cg13859478) in the gene *CANX* was significantly associated with its protein abundances but not with its mRNA expression levels. Figure 3 shows the heatmap of the five omics data types of the gene *CANX* in the 69 overlapping samples. Lastly, 89 genes (93 methylation sites) were only associated with the corresponding mRNA levels but not global protein or phosphoprotein abundances.

**Figure 3:**
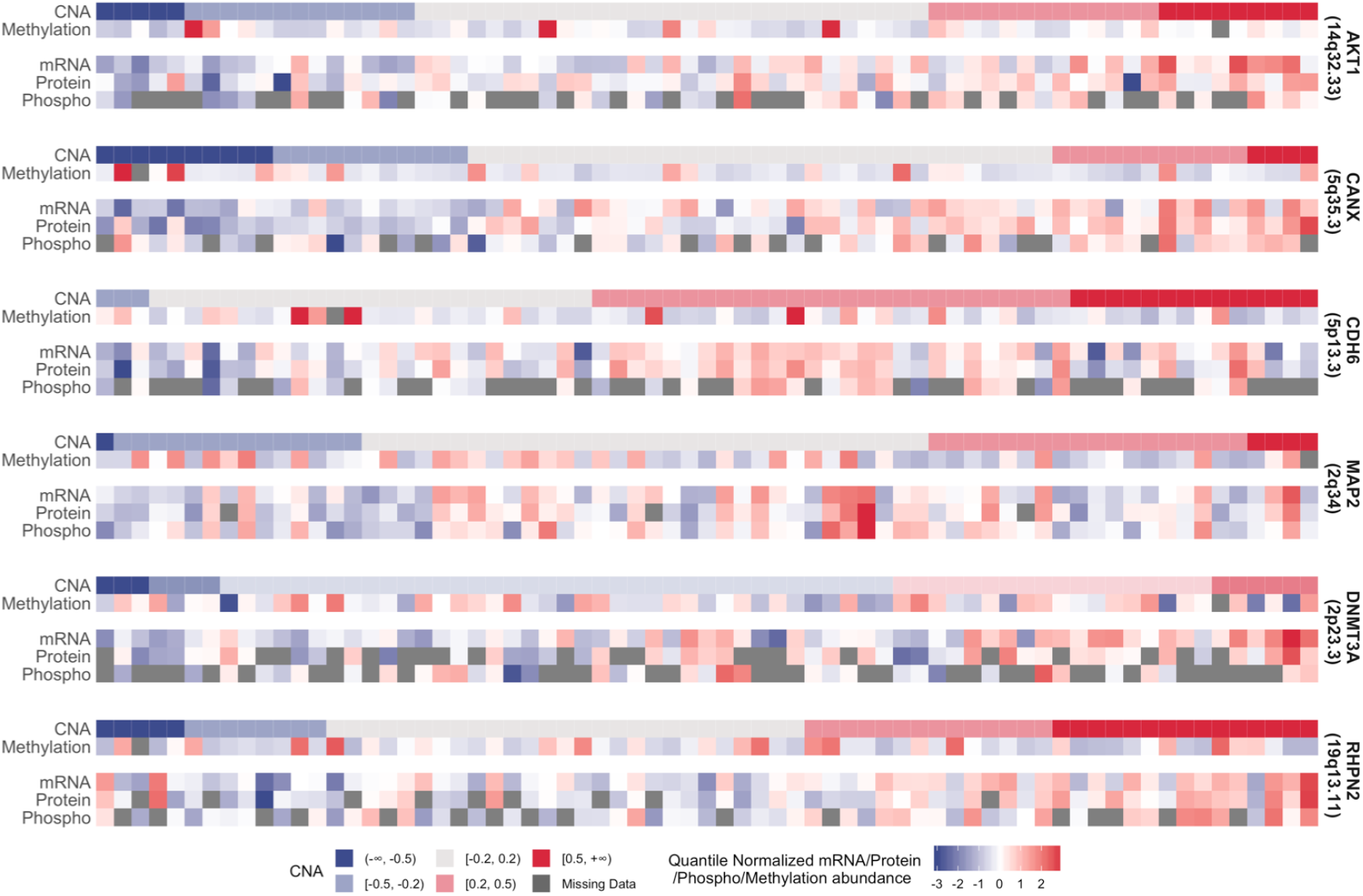
The heatmap of key identified genes among the 69 overlapping samples in which all of the five data types have been measured. *AKT1* is the key hub of the cascade CNA network; the methylation of CANX plays a regulatory role on protein abundance, without mRNA reflection; the genes *DNMT3A, RHPN2, MAP2* and *CDH6* have cascading CNA effects and methylations effects on cis-mRNAs and proteins.

Figure 4 demonstrates the genome distribution of all CNAs and methylations with significant cis-associations. Two cascade CNAs hotspots are identified on 14q and 19p with at least five genes within 10MB showing cascade CNA cis-regulations. The 14q hotspot sits in 14q32.31-32.33 and contains six cascade CNAs (AKT1, MTA1, DYNC1H1, CDC42BPB, EIF5 and PPP2R5C), while the 19p hotspot sits in 19p13.2-13.11, harboring seven cascade CNAs (PRDX2, RAB8A, BRD4, GIPC1, MYO9B, GATAD2A, FKBP8). Detailed depictions of all omics data related to these genes within the two hotspot regions are presented in Supplementary Figures S3 and S4.

**Figure 4:**
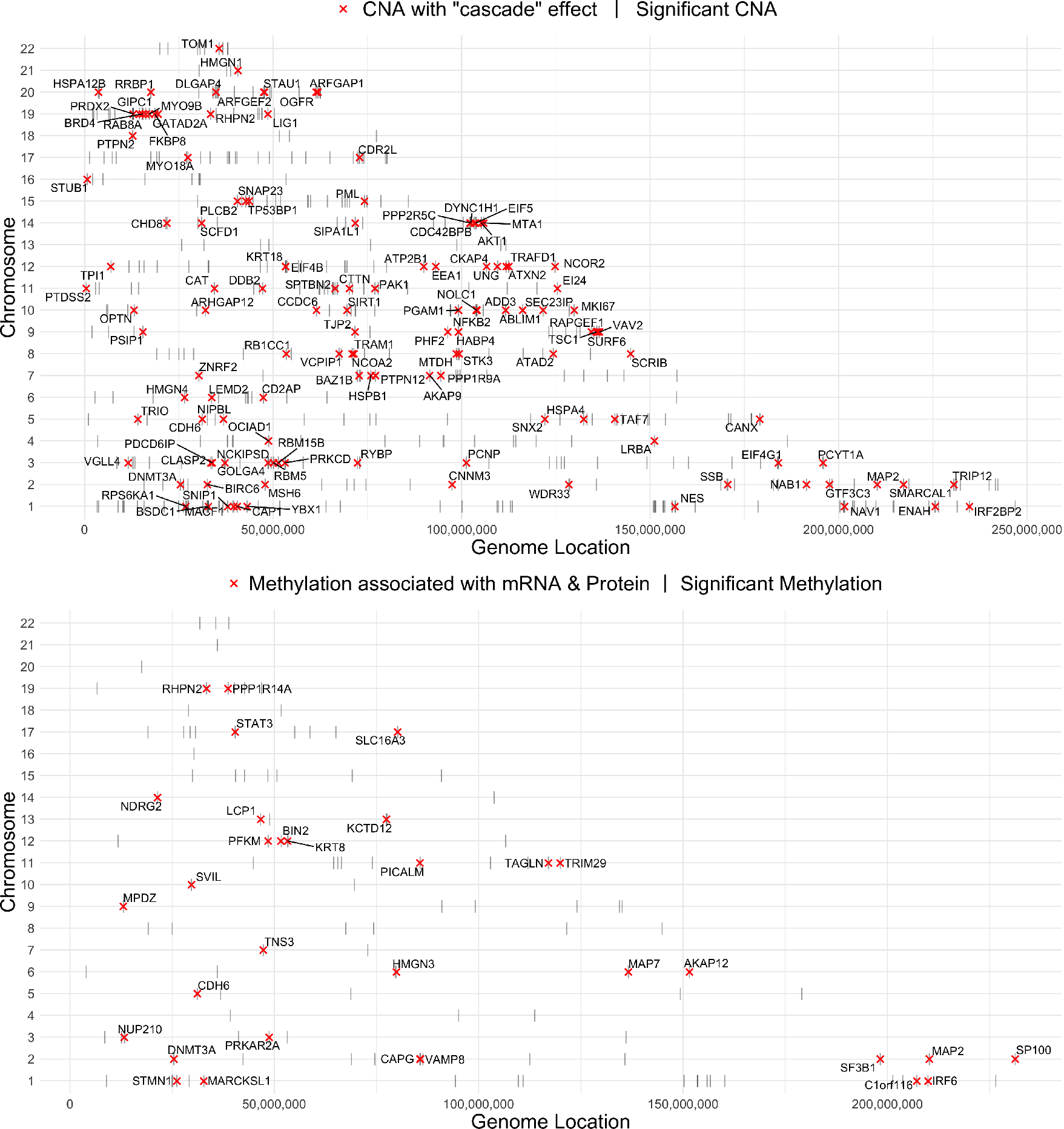
The genome distribution of identified CNAs and DNA methylations.

### Comparison with a conventional method and a power study

We next compared the power and false discovery control of iProFun to existing approaches. iProFun employs empirical assessment of FDR instead of using commonly used FDR control procedures for the following reasons: the q-value procedure [33] and Benjamini and Hochberg (BH) procedure [34] assume independence among multiple tests, and the revised Benjamini and Yekutieli (BY) approach [35] allows only weak dependency among different tests. Since there are often unknown yet potentially high correlations among nearby genes, none of them holds valid FDR control (Supplementary Figure S6). Therefore, empirical assessment through permutation provides more accurate assessment and should be applied for error control.

With the empirical error control, Figure 5(a) plots the number of genes discovered under our iProFun method and a conventional approach that separately considers each molecular quantitative trait in the analysis. The conventional approach only uses samples that contain the molecular quantitative trait of interest for regression, adjusting for the same covariates as our integrative analysis. Using the same FDR criteria (empirical mean FDR < 10%), iProFun identifies many more genes than separate analyses, by borrowing information across data types. We observed higher power gain in the data types with smaller sample sizes, such as protein and phosphoprotein abundances. We observed substantial power improvement in identifying global protein QT CNA (pQTC) (from 430 to 501 genes), phosphoprotein QT CNA (phQTC) (from 0 to 129 genes) and global protein QT methylation (pQTM) (from 0 to 261). Neither of the approaches was powerful enough to identify any significant phosphoprotein QT methylation (phQTM). Our integrative analysis pipeline greatly boosts study power for gene identification, especially for the data types with small initial sample sizes.

In Figure 5(b) we compared the number of identified genes using all samples available to us through TCGA-CPTAC to the number identified using a subset of samples. In particular, while both approaches used the same sets of samples for mRNA and phosphoprosites, the subset approach narrowed our analysis of protein abundances to the 69 samples that also have measurements on phophosites. Such exercises serve two purposes: (1) with identical samples in global and phosphoprotein data, we can compare the signal strengths of these two data types, and (2) we can evaluate the power gain for all data types in the integrative analysis via iProFun by adding samples to one data type. The full bar represents the number of genes identified with FDR<10% in full samples and sub-samples (same as Figure 5(a)), while the intensified proportion also imposes posterior probabilities > 75% to require high confidence of the discoveries. Under both criteria, having more protein samples integrated via iProFun will greatly boost power for pQTC and pQTM identification, while it may also increase identifications for other data types.

**Figure 5:**
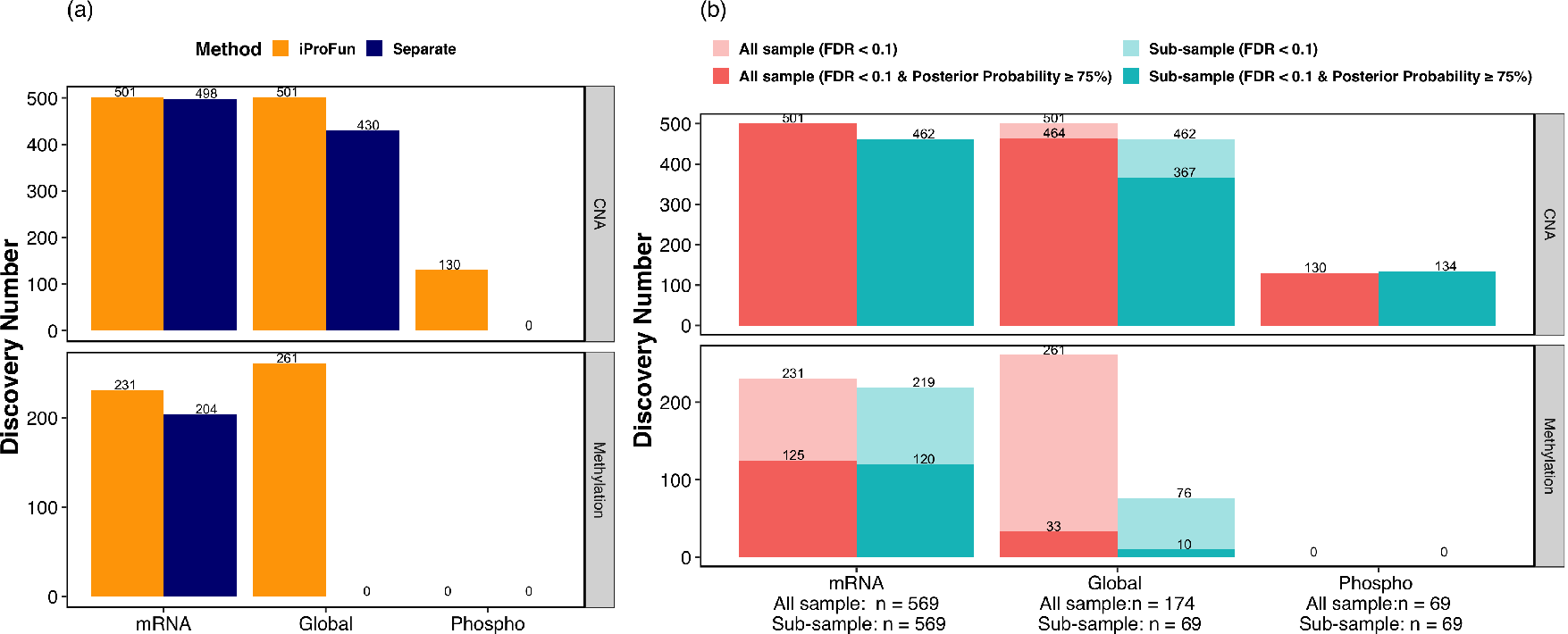
(a) Comparison between integrative and separate analyses on the number of identified genes (FDR<10%). (b) Comparison of the number of identified genes using all samples available and a subset of data with 69 samples having global protein abundance.

## DISCUSSION

In this study, we introduced a novel integrative analysis tool, iProFun, to more powerfully recognize proteogenomic functional traits altered by CNA and DNA methylation by jointly modeling CNA, epigenome, transcriptome, global proteome and phospho-proteome data. This integrative solution boosts power for detecting significant cis-associations by borrowing information across different omics data types.

We applied iProFun to the ovarian HGSC tumor data from TCGA and CPTAC. We chose this cancer target data set primarily for two reasons. First, HGSC is the leading cause of gynecologic cancer death in the US [23]. Most women will present with advanced-stage disease and then undergo cytoreductive surgery followed by combination platinum-based chemotherapy. Despite initial response to primary therapy, the majority of women will experience disease recurrence and ultimately die of their disease within five years [24]. Thus, new treatments and an improved understanding of the biologic basis of this cancer are desperately required. Second, in the TCGA-CPTAC HGSC project, for the first time, large-scale deep proteome, especially phospho-proteome, analyses were employed to study HGSC by research teams from CPTAC, a national effort to accelerate the understanding of the molecular basis of cancer. This provides us an unprecedented opportunity to integrate five different types of omics profiles of the same set of tumors to better understand the impact of DNA alterations on all mRNA, protein and phospho protein activities in HGSC.

Using iProFun we identified a large number of genes whose CNAs and DNA methylation significantly affected the genes’ mRNA, protein and/or phosphoprotein activities. Specifically, iProFun identified 130 CNAs that impact all levels of molecular QTs, i.e. mRNA, global and phosphoprotein abundances; our definition of “cascade” cis-effects. This set should be enriched for biologically relevant cancer genes, as CNAs with preserved functional consequences are more likely to be cancer drivers. A network analysis of these 130 genes using the STRING database directed our attention to the gene *AKT1*, a key hub in the network, which interacts with many other cascade CNA genes (Supplementary Figure S5). The gene *AKT1* is an effector in the PI3K/RAS pathway, which is deregulated in nearly half of all HGSC cases [26] and down-regulation of its phosphoprotein was found to be associated with poor survival outcomes in the original TCGA-CPTAC ovarian study [16]. While the impact of AKT1 copy number alteration was not discussed in the previous ovarian studies [16], iProFun results suggest that down-regulation of AKT1 phosphopeptides are associated with DNA copy number losses of the gene. This potentially therapeutically-targetable association/pathway, as well as the other 129 cascade CNA cis-associations, however, would not have been detected if only CNA phosphoproteomics data alone would have been analyzed due to the challenge of high-dimension-low-sample-size in such investigations (Figure 5).

Another interesting finding by iProFun is that one methylation site (cg13859478) of the gene *CANX* revealed significant cis-associations with its global protein abundances but not its mRNA expression levels. To the best of our knowledge, this is the first association of CANX with ovarian cancer. The gene *CANX* is a member of calnexin family of molecular chaperones. Its encoded protein is a calcium-binding, endoplasmic reticulum (ER)-associated protein that interacts transiently with newly synthesized N-linked glycoproteins, facilitating protein folding and assembly. Over-expression of CANX has been identified as a favourable prognostic marker in colorectal cancer but unfavourable prognostic marker in thyroid cancer [36]. This cancer-type specific effect of expression in different cancers might be partially explained by the iProFun results that methylations of CANX were not associated with its mRNA levels. Further analyses on global protein and phophoprotein across cancers might shed light on explaining the contrasting effects previously noted by others.

Finally, there were a set of four genes that were overlapping in the 130 cascading CNA genes and 32 genes with methylations associated with both mRNA and protein levels. The four genes were *DNMT3A, CDH6, RHPN2* and *MAP2*. All four have been previously shown to play different and important roles in cancer but only DNMT3A and CDH6 have been previously linked to ovarian cancer. *DNMT3A* encodes a DNA methyltransferase. CpG methylation is an epigenetic modification that is important for embryonic development, imprinting, and X-chromosome inactivation. The molecular QTs of this gene are associated with both its CNA and DNA methylation. The dysregulation of the gene *DNMT3A* is critical in the development of certain cancers, including pancreatic, endometrial, liver and renal cancer [37, 38, 36]. In ovarian cancer, associations between DNMT3A expression and a number of different microRNAs have been shown to have biologic effects on cell growth and outcome [39, 40, 41]. *CDH6* encodes a member of the cadherin superfamily. Cadherins are calcium-dependent cell adhesion proteins that play critical roles in cell differentiation and morphogenesis. *CHD6* has been shown to be both highly differentially expressed in ovarian cancer and, taking advantage of its surface expression, demonstrated to be a unique therapeutic target for antibody-drug conjugates [42].

*RHPN2* encodes a member of the rhophilin family of Ras-homologous (Rho)-GTPase binding proteins. It may limit stress fiber formation and/or increase the turnover of F-actin structures, and has been identified as a prognostic marker in renal and head and neck cancer [37, 38, 36]. Amplification of the RHPN2 locus has been associated with a poorer outcome in glioma and ectopic overexpression of RHPN2 in cell lines promoted an invasive phenotype [43]. Finally, *MAP2* encodes a protein that belongs to the microtubule-associated protein family, and has been identified as a prognostic marker in endometrial cancer [37, 38, 36].

As already noted, in part, one of our reasons for targeting ovarian cancer in our studies was the current relatively bleak landscape for novel therapies. Thus, it is notable that our analyses have identified a number of therapeutic candidates. All of *AKT1, DNMT3A* and *MAP2* are druggable genes with approved drugs already on the market with indications for other tumors [44]. Our results suggest that integrating multiple-omics data to screen for genes whose DNA alterations have significant impact on functional molecular traits can be a very effective strategy to nominate candidate genes contributing to the disease. While we believe that CNAs and DNA methylations, which play key roles in disease etiology of cancer, should preserve functional consequences, not all genetic alteration events with functional impacts are disease relevant. Thus, after iProFun is performed to nominate disease relevant genes, further investigation leveraging additional information, such as gene-gene interaction network and patient outcome data based analyses could further help to pinpoint the most promising candidates.

Beyond the current studies, while we focus on the functional regulations of somatic CNA and DNA methylation, iProFun provides a general framework that can be easily extended to a wide range of applications. For example, by considering the same molecular quantitative trait from three subtypes as if it were from three omics data types with no overlapping samples, we could identify CNAs and methylations whose functional consequences are shared across subtypes versus CNAs and methylations whose functional consequences are unique to one or some of the subtypes. We could also extend the tool to germline variations, adding additional molecular QTs, and/or *trans* associations analysis. To provide a balanced perspective, we also note potential limitations of our analysis. First, iProFun is based on a linear regression framework and calculated the posterior probabilities using *t*-distributions, which might not be directly applicable to analyses that follow other distributions (e.g. *χ*^2^, *F* or Uniform distributions). Second, iProFun requires a relatively large number of genomic features, such as CNAs and DNA methylations, to estimate the density under the alternative distribution and robustly calculate the posterior probabilities. In cases where only a few genes are quantified, iProFun might not provide optimal results. Future studies could expand iProFun to incorporate more association analysis tools (e.g. provide a p-value based algorithm) to overcome these limitations.

Software implementing the proposed iProFun, as well as CNA, DNA methylation, mRNA, global and phosphoprotein data used for this analysis, are available on Github, https://github.com/songxiaoyu/iProFun.

## Supplementary Materials

**Figure S1:**
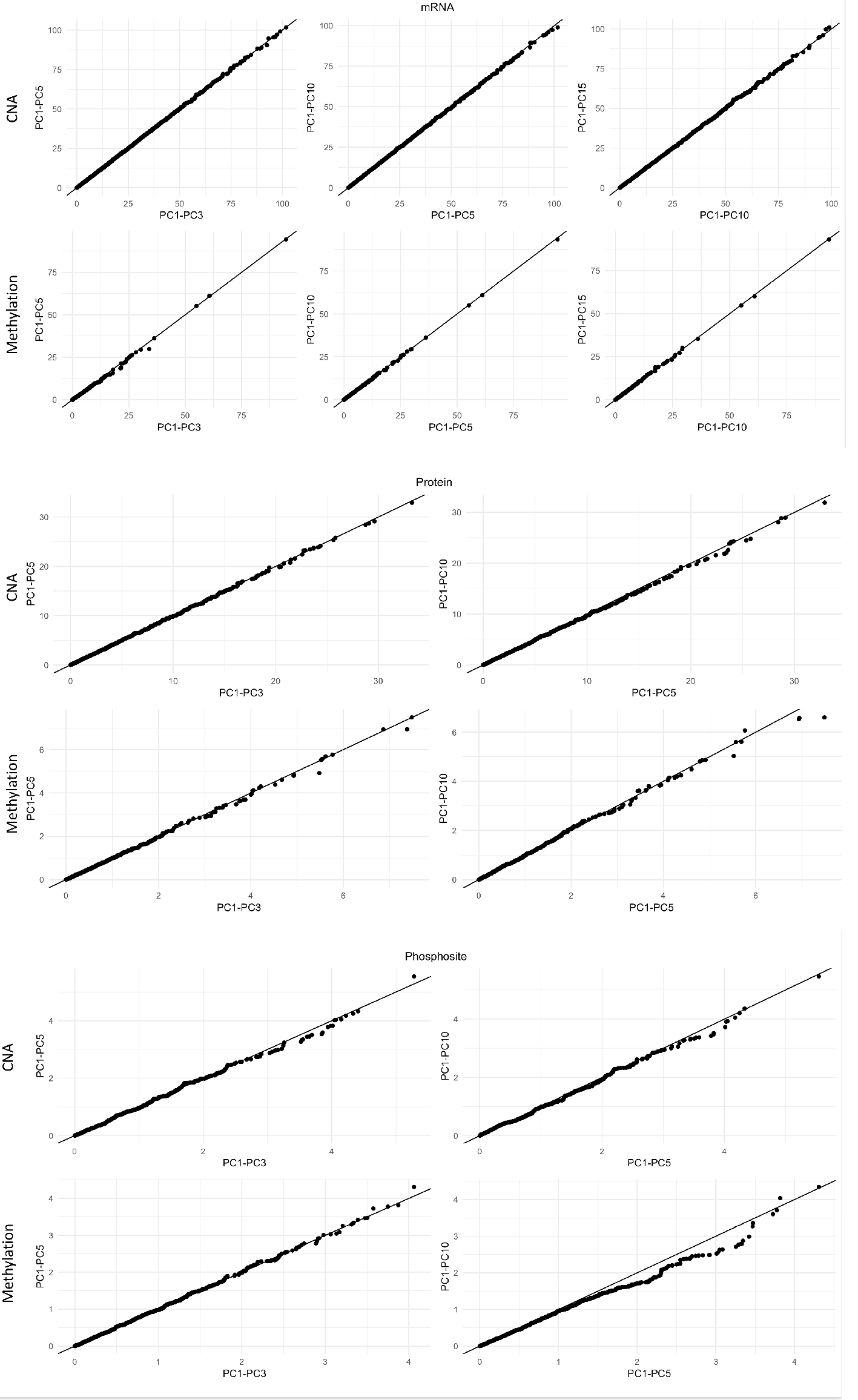
The QQ-plot of regression –*1og*_10_ (p-values) adjusting for different principal components (PCs). Top 3 PCs were selected for all three molecular quantitative traits of mRNA, global protein and phosphoprotein to account for population stratification and other major confounding factors.

**Figure S2:**
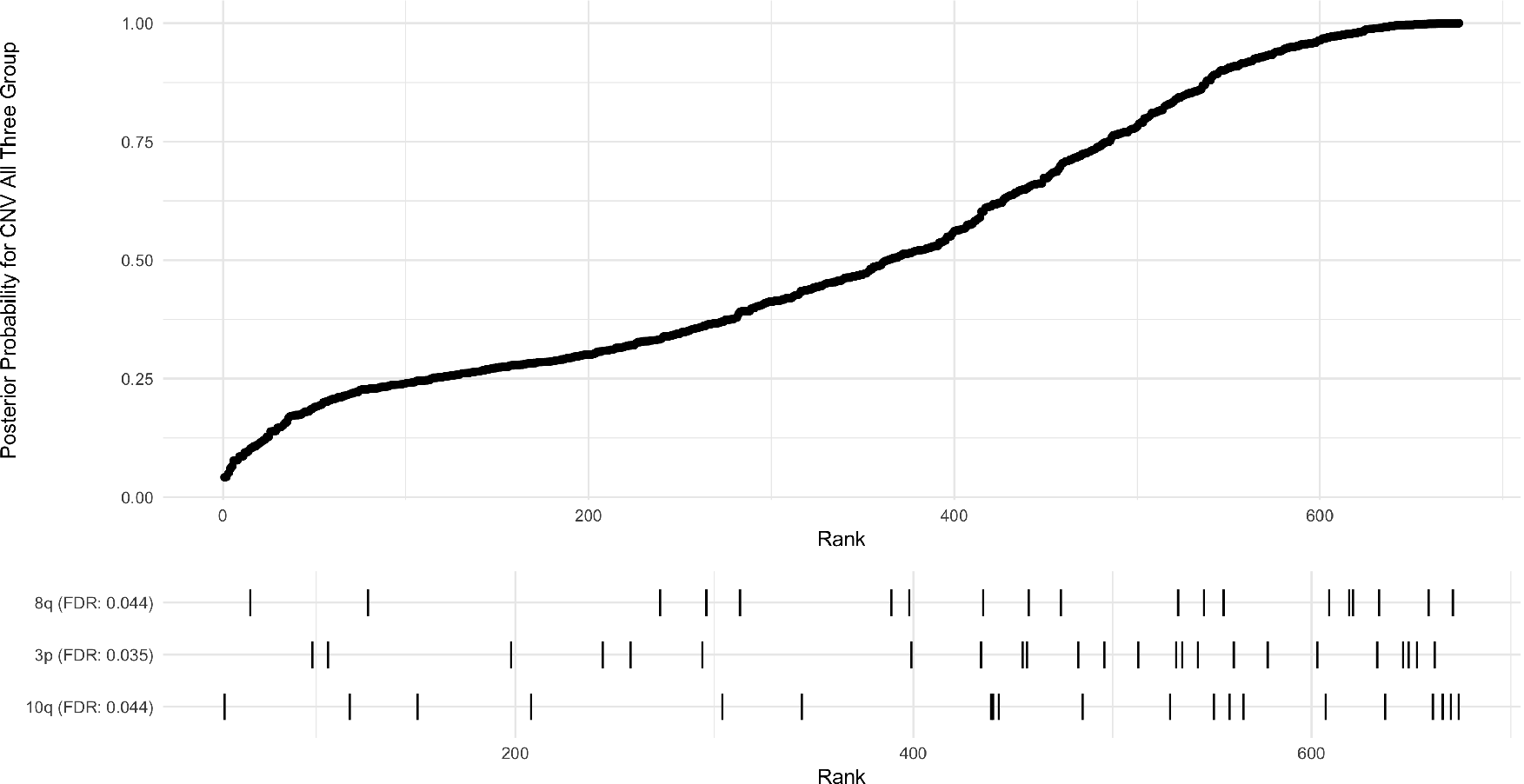
The chromosome arm imbalance enrichment of CNAs and methylations with “cascade” events. The cascade CNAs were significantly enriched in arm 3p (FDR=0.034), as well as 8q and 10q (both FDR=0.043).

**Figure S3:**
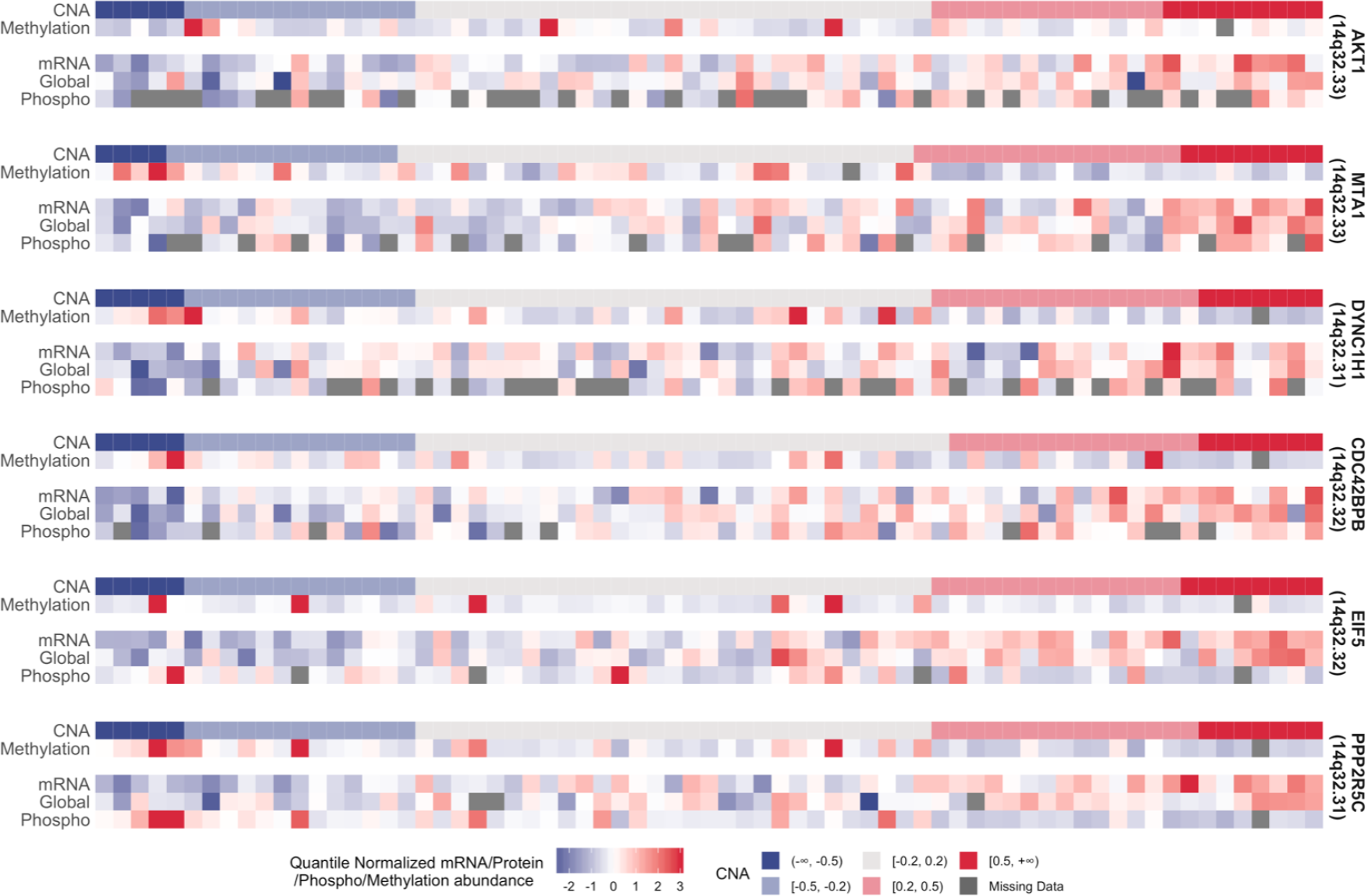
Heatmap of genes from 14q32.31-32.33 CNA key hub (*AKT1, MTA1, DYNC1H1, CDC42BPB, EIF5, PPP2R5C*)

**Figure S4:**
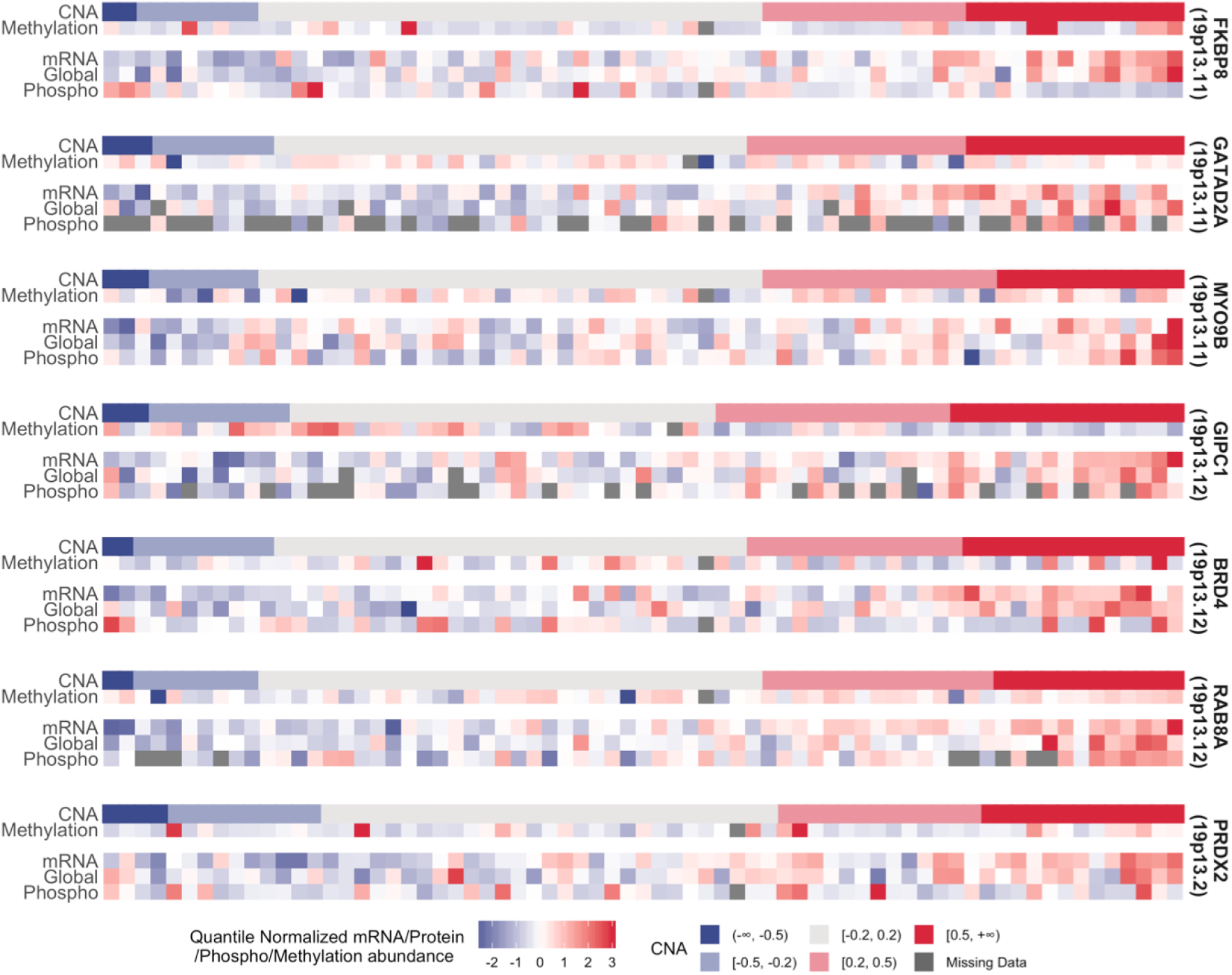
Heatmap of genes from 19p13.2-13.11 CNA key hub (*FKBP8, GATAD2A, MYO9B, GIPC1, BRD4, RAB8A, PRDX2*

**Figure S5:**
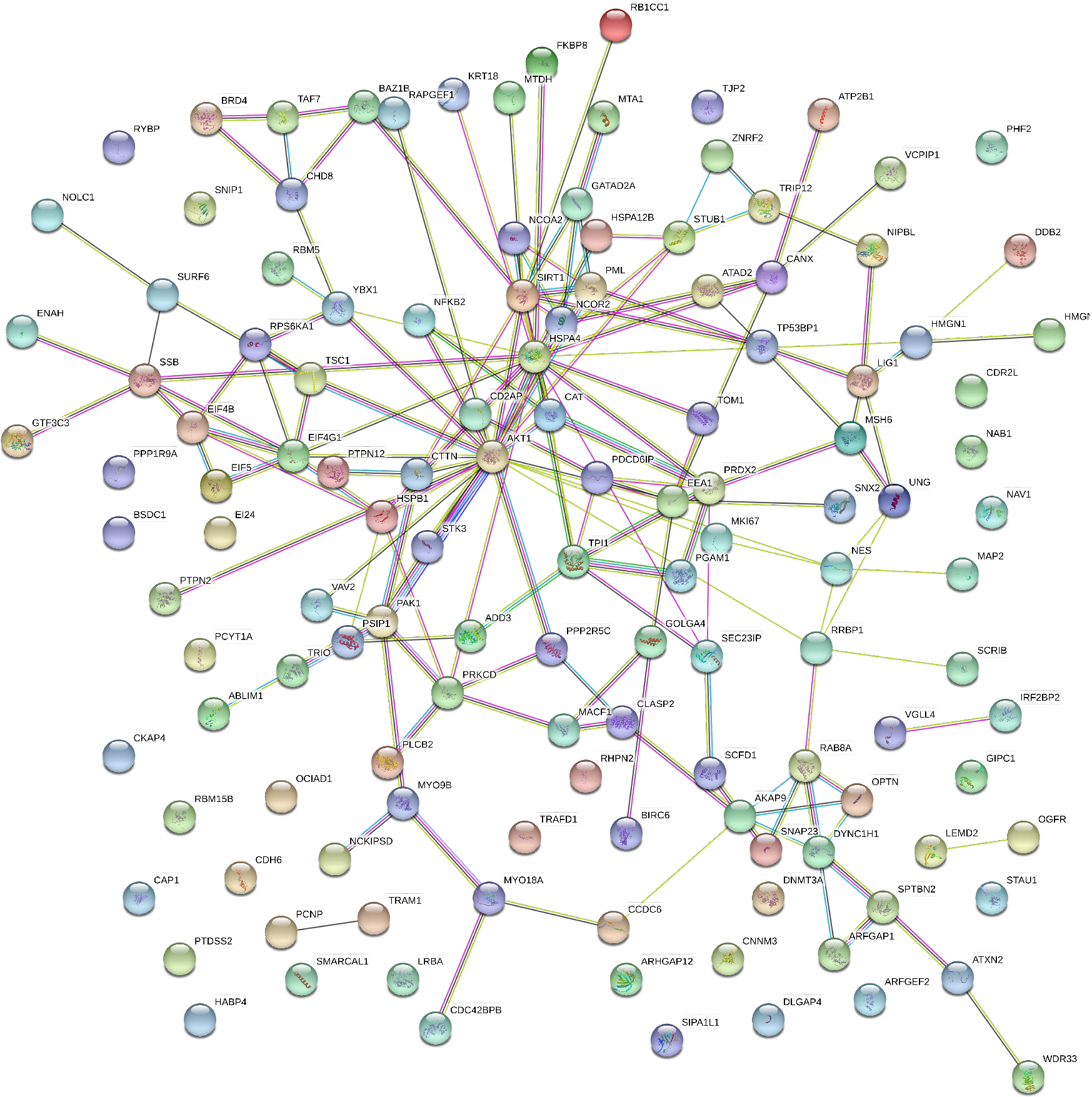
String Network of 130 CNAs with “cascading” effects. Gene *AKT1* is a key hub node.

**Figure S6:**
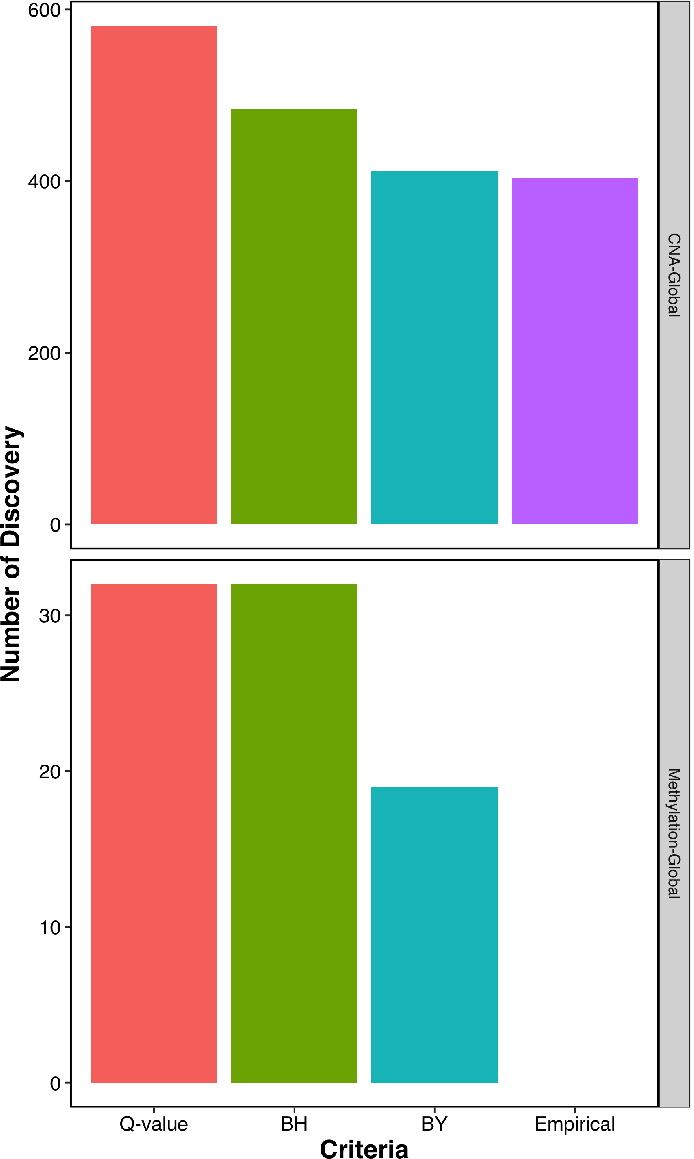
Comparison of traditional FDR control methods with empirical assessment that we applied in iProFun pipeline. The commonly used q-value procedure and Benjamini and Hochberg (BH) assume independence among multiple tests, and the revised Benjamini and Yekutieli (BY) approach allows weak dependency among different tests. Since there are often unknown yet potentially high correlations among nearby genes, none of them holds valid FDR control. Therefore, the assumptions of commonly used FDR approaches are violated in this application, and we have employed empirical assessment through permutation in our iProFun pipeline.

